# Anticipatory reinstatement of expected perceptual events during visual sequence learning

**DOI:** 10.1101/2020.11.28.402123

**Authors:** Mehdi Senoussi, Rufin VanRullen, Leila Reddy

## Abstract

Being able to predict future events in learned sequences is a fundamental cognitive ability. Successful behavior requires the brain to not only anticipate an upcoming event, but to also continue to keep track of the sequence in case of eventual disruptions, (e.g., when a predicted event does not occur). However, the precise neural mechanisms supporting such processes remain unknown. Here, using multivariate pattern classification based on electroencephalography (EEG) activity and time-frequency amplitude, we show that the visual system represents upcoming expected stimuli during a sequence-learning task. Stimulus-evoked neural representations were reinstated prior to expected stimulus onset, and when an anticipated stimulus was unexpectedly withheld, suggesting proactive reinstatement of sensory templates. Importantly, stimulus representation of the absent stimulus co-occurred with an emerging representation of the following stimulus in the sequence, showing that the brain actively maintained sequence order even when the sequence was perturbed. Finally, selective activity was evident in the alpha-beta band (9-20 Hz) amplitude topographies, confirming the role of alpha-beta oscillations in carrying information about the nature of sensory expectations. These results show that the brain dynamically implements anticipatory mechanisms that reinstate sensory representations, and that allow us to make predictions about events further in the future.

## Introduction

The brain makes predictions about upcoming events based on regularities that have been learned in the past. This ability to extract and learn associations between events allows us to anticipate the future, and prepare appropriate actions and decisions accordingly. In the real-world however, expectations are often violated. For instance, one may have learned the order of traffic signs on a familiar route, but may occasionally find one of these signs missing. In these situations, when an event sequence is momentarily and occasionally disrupted, a crucial requirement of predictive neural mechanisms is to be able to deal with the missing information: the brain must be able to keep track of where it was in the sequence, and continue to make predictions about what will happen next.

Theoretical accounts (de Lange et al., 2018) predict that prior information about upcoming events biases neural representations. Several studies have shown that an expected stimulus induces sensory representations in visual and auditory cortex (Demarchi, 2019; deLange, 2018; Miyashita, 1988; Kok et al., 2012a, 2017), and evokes anticipatory activity in higher-level structures (Schapiro et al., 2012; Davachi and DuBrow, 2015; Reddy et al., 2015). Less clear is how the brain copes with disrupted expectations. At the perceptual level, when expectations are invalid, the speed of stimulus detection may be delayed (de Lange et al., 2018). However, it is unknown how the prediction of events further into the future is affected when a current expectation is violated (e.g., when an expected event does not happen).

In the current study, we investigated whether the brain actively and dynamically predicts future events in a sequence before they occur. Moreover, we asked what happens when the sequence is disrupted and an anticipated event does not occur. Does the brain still actively maintain its position in the sequence, and can it predict what comes next? Finally, we tested whether the neural signatures of expectation were supported by specific oscillatory components.

Using multivariate, time-resolved electroencephalography (EEG) decoding and a visual sequence learning task, we observed anticipatory reinstatement of stimulus-evoked neural patterns before each stimulus appeared. Furthermore, these anticipatory neural activity patterns for future stimuli were not erased when the sequence was disrupted and the next stimulus did not appear. They persisted even after incoming sensory information indicated the absence of the expected stimulus, concurrently with the emergence of the neural patterns of the next stimulus in the sequence. These results suggest dynamic mechanisms that activate neural representations for upcoming stimuli ahead of time, and keep track of sequence information even when expectations are violated. These anticipatory mechanisms might allow the brain to make predictions about future events, and to cope with missing information during sequence maintenance.

## Results

### Task and behavioral results

Fifteen participants were presented with a sequence of 6 images in a fixed order (Reddy et al., 2015) while their brain activity was recorded with EEG. The image sequence was repeated 480 times (i.e. 6 images*480 iterations of the sequence = 2880 trials). On “stim” trials (45% of trials), each image was presented for 1s. On these trials, the participants’ task was to learn the order of images in the sequence. Participants were tested on how well they learned the sequence order on a random 10% of trials (“test trials”). On these “test” trials the sequence stopped, and participants were presented with two choice stimuli and asked to report which of the two was the next one in the sequence. After their report, the sequence resumed. On the remaining 45% of trials (“catch trials”, occurring randomly), the expected stimulus was omitted and replaced by a gray square for the 1s duration of the trial. In order to be able to perform successfully on the subsequent test trials, this design required participants to actively maintain sequence information on catch trials. Each trial (stim/catch/test) was preceded and followed by an inter-stimulus interval (ISI) of 0.5s (SI Appendix, Fig. S1A).

Accuracy on the 2-alternative forced choice (AFC) task performed during test trials was above chance in all blocks (one-sample t-test: t(14) > 8, P < 10^-6^), but increased across the 8 blocks performed by each participant (one-way, random-effects ANOVA: F(7, 120) = 5.55, P < 5.10^-5^). A post-hoc Tukey’s HSD showed that only the first block was significantly different from the others (P < 0.05). Participants therefore rapidly learned the stimulus sequence by the end of the first block of the experiment.

### Decoding of stimulus identity in the sequence

We first asked if EEG activity patterns carried stimulus-specific information for the six stimuli in the sequence. A multivariate six-way pattern classifier was trained on the EEG responses to discriminate between the six visual stimuli on the “stim” trials, when the stimuli were visually presented to the participants, and a temporal generalization approach (King and Dehaene, 2014) was used to evaluate how stimulus-specific information evolved over the time course of the trial. Decoding performance of these “stim-classifiers” was successful and highly above chance (peak accuracy: 64.4%/3.25 (mean/s.e.m); P < 10^-4^; SI Appendix, Fig. S1B) indicating that EEG response patterns uniquely represented each of the six stimuli in the sequence. Activation patterns, reflecting the discriminative topography used by the classifier to distinguish the different classes (see *Materials and Methods*), revealed a sequence of right-occipital, occipito-frontal and parietal topographical distributions across the stim trial time course.

### Selective neural activity for anticipated events

Do brain activity patterns also uniquely encode anticipated events in the absence of visual input? To address this question, we first examined the ISI periods before the onset of each trial. Because the sequence was repetitive and the image order predictable, in each ISI period participants could anticipate the next upcoming stimulus of the sequence. There were therefore six possible ISI period events, based on which image they preceded. In this analysis we asked if these six ISI events could be distinguished from each other based on EEG activity patterns. Because the ISI periods were not predictive about whether the subsequent trial would be a stim, catch or test trial, all ISI periods were pooled together in this analysis.

Since no visual stimulus was presented during these ISI periods, significant decoding performance observed in the ISI periods would reflect a unique anticipatory preparation for the upcoming, expected event. As above, we used temporal generalization to distinguish the six ISI events. The different ISI events were discriminable from each other, starting around 250ms-300ms prior to trial onset (P < 0.0001, Fig. 2A). Decoding performance exhibited a ramping selectivity for the upcoming event, in the absence of any visual input, suggesting anticipatory preparation for the immediately following event. Activation patterns revealed a succession of right-occipital and occipito-frontal topographical distributions across the ISI time course (Fig. 2A), similar to those observed at the beginning of stim trials (SI Appendix, Fig. S1B).

**Figure 1.**
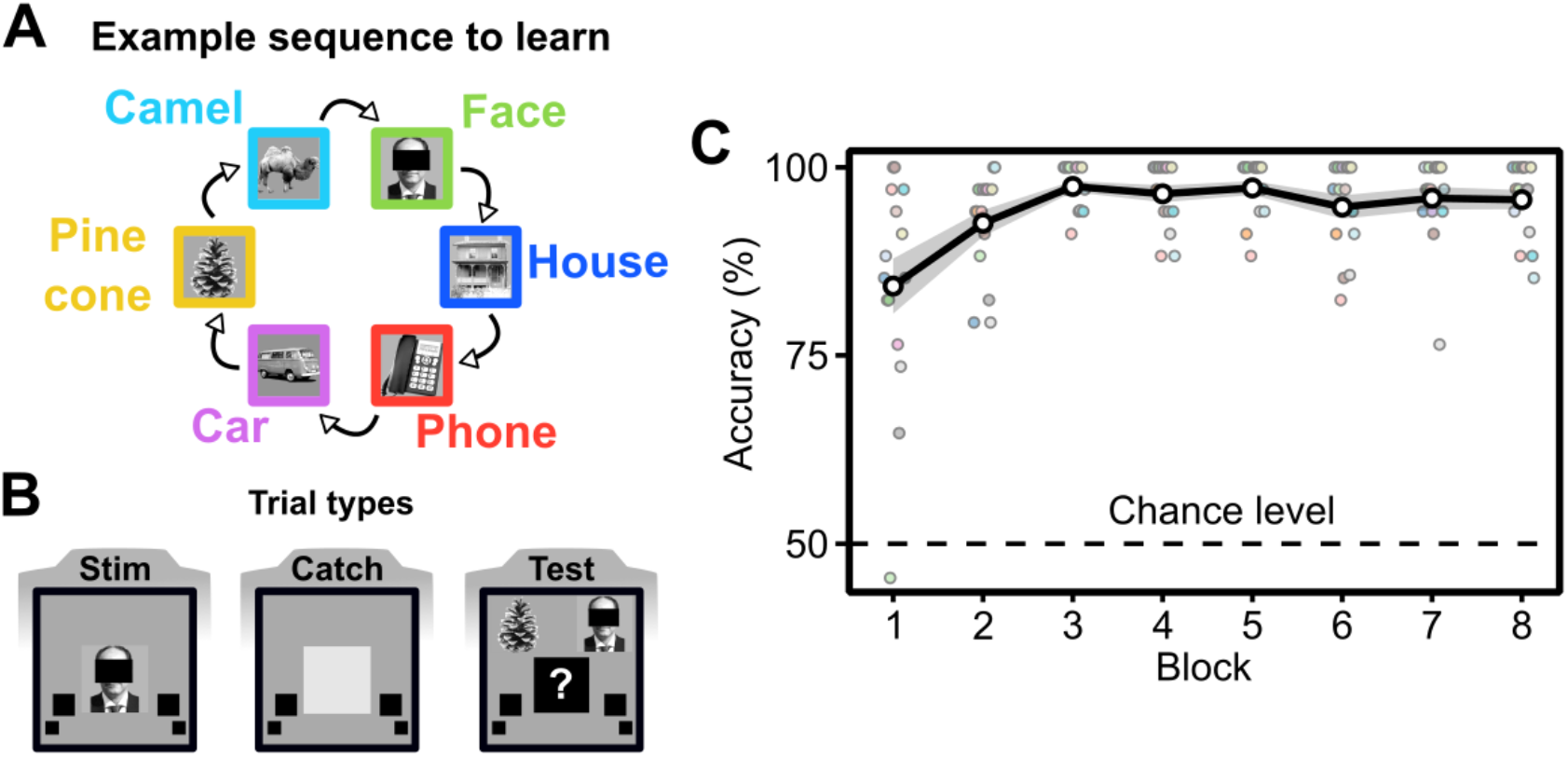
Experimental protocol and behavioral results. **A.** Fifteen participants were asked to learn the order of a sequence of six stimuli. **B.** The experiment consisted of three trial types. On stim trials the expected stimulus was shown. On catch trials, the expected stimulus was replaced by a gray square. On test trials, participants were presented with two choice stimuli and had to report which of the two was the next one in the sequence. Each stimulus occurred for 1s and was preceded by an inter-stimulus interval (ISI) of 0.5s. **C.** Accuracy on the 2-AFC task during test trials. Participants achieved more than 80% accuracy on average in the first block. White dots represent the average across participants, gray shading around the curve represents standard error of the mean and each small dot represents a single participant’s accuracy.

**Figure 2.**
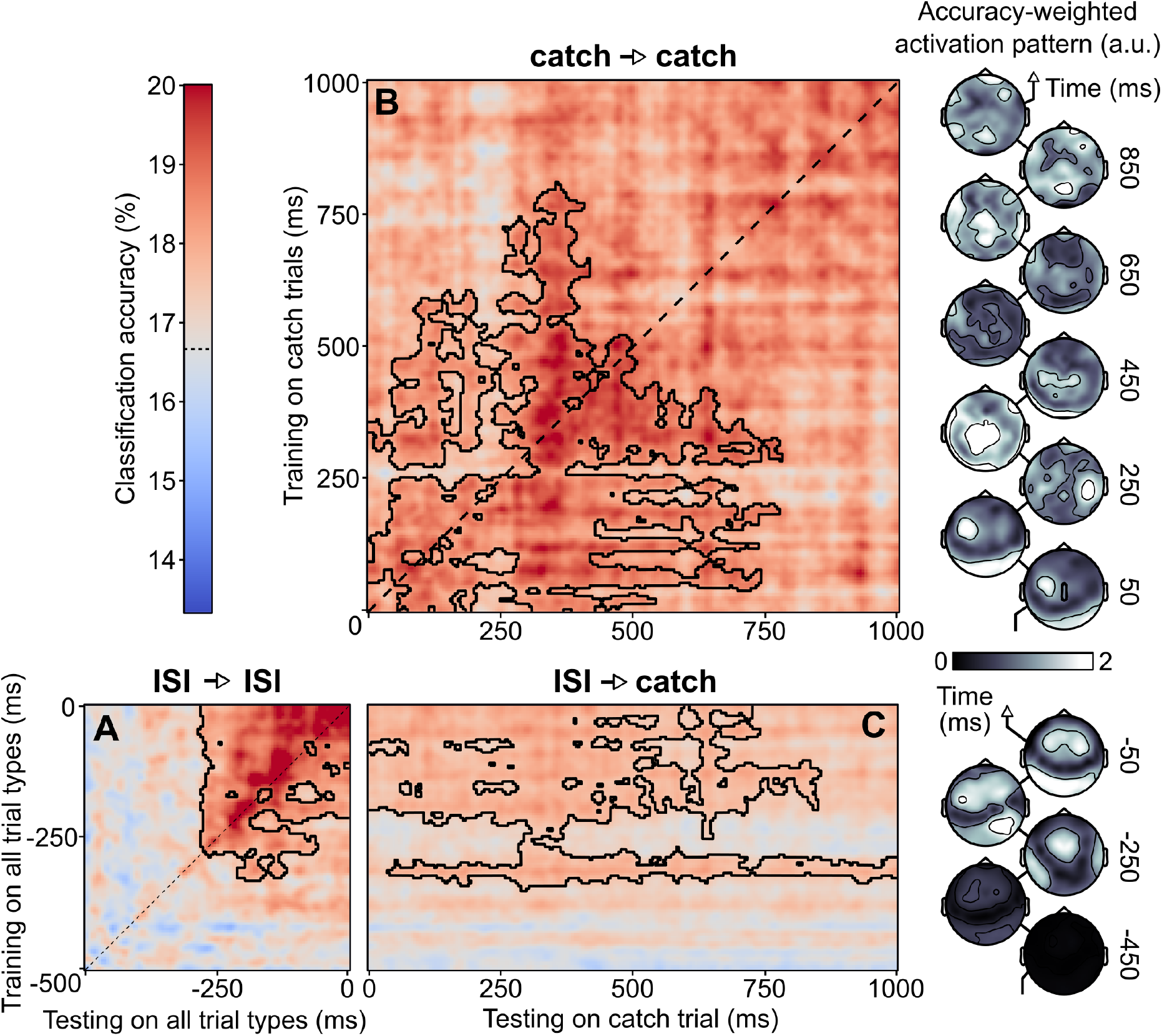
Classification performance during the ISI periods and the catch trials. **A.** Temporal generalization on ISI periods. A classifier was trained at each time point (vertical axis: training time) and tested on all time points (horizontal axis: testing time). The black dashed line represents the diagonal of the temporal generalization matrix, i.e. classifiers trained and tested on the same time point. Black contours indicate a cluster of significant above-chance classification accuracy (P = 0.0001). **B.** Temporal generalization on catch trials (0 to 1000ms with respect to trial onset). Same arrangement as in A. Black contours indicate a cluster of significant above-chance classification accuracy (P < 0.02). Topographies on the right side of the map represent activation patterns weighted by decoding performance (see *Materials and Methods*) for classifiers trained on catch trials. **C.** Cross-temporal generalization in which classifiers were trained on data from the ISI period and tested on catch trials. Black contours indicate a cluster of significant above-chance classification accuracy (P < 0.002). Topographies on the right side of the map represent activation patterns weighted by decoding performance (see *Materials and Methods*) for classifiers trained on ISI periods.

We next explored whether selective patterns of neural activity can be induced purely as a result of top-down expectation signals, (i.e., in the “catch” trials when stimuli were expected, but omitted (Fig. 2B)). As above, we used temporal generalization to distinguish the six catch events from each other (i.e., one catch event per stimulus type) across the entire catch trial epoch (from 0ms to 1000ms relative to trial onset). The catch events could be discriminated from each other over almost the entire duration of the catch trial window, starting from stimulus onset to around 800ms post stimulus onset (P < 0.02, Fig. 2B). Since in the catch trials all stimulus information was replaced by a gray square, this successful decoding of catch trial activity necessarily reflects a top-down representation of the expected event. Activation patterns revealed occipital and mostly parietal topographical distributions across the catch trial time course (Fig. 2B).

Anticipatory neural activity patterns elicited during the ISI periods were also informative about the subsequent catch trials. A cross-temporal generalization analysis in which classifiers were trained on data from only the ISI period and tested on data from the post-trial onset period of catch trials revealed that the selective neural patterns found during the ISI period generalized to the post-trial onset period (cluster test, P < 0.002), (Fig. 2C). These results indicate that predictive patterns of neural activity elicited prior to the onset of an expected event persisted during the event itself.

These results thus indicate that unique neural activity patterns are instantiated in anticipation of a predicted event, and importantly, that these patterns are maintained throughout the duration of the expected event, even when it ultimately fails to occur. These activity patterns thus might be critical markers that allow the brain to keep track of the progress of the sequence when the sequence is disrupted. However, do these “markers” only signal sequence position, or are they also informative about the identity of the upcoming and/or missing stimuli?

### Neural activity patterns for anticipated events are informative about stimulus identity

To determine whether the brain activity patterns that we observed above for anticipated stimuli are merely generic signals marking the progress of the sequence, or whether they actually carry information about the identity of one or more upcoming stimuli, we performed a cross-temporal generalization analysis. In this analysis, a multivariate “stim-classifier” was trained on EEG data to predict stimulus identity. This stim-classifier was then tested on 1) the ISI periods before trial onset, and 2) the post-trial onset periods of catch trials (i.e., in the absence of stimulus input). This approach allowed us to extract neural activity patterns that uniquely identify the different stimuli, and use them to test whether these same patterns were evoked by purely predictive mechanisms when no stimulus was presented. In other words, are stimulus-evoked neural representations reinstated when the same stimuli are expected?

More precisely, the stim-classifier was trained to discriminate between the 6 images, at each time point of “stim” trials (150ms after stimulus onset, see *Materials and Methods*). It was then used to decode stimulus identity when stimuli were expected but not shown (i.e., during the ISI periods and catch trials). For example, consider a segment of the sequence consisting of three stimuli: face-house-phone, and let us label them stimuli N-1, N, and N+1 respectively (SI Appendix, Fig. S2B). Let us assume that the sequence is currently at stimulus N (i.e., the house), but that on this iteration of the sequence, this trial is a catch trial i.e., the house was expected, but it actually did not materialize (“catch house trial”). If the stim-classifier is tested on data recorded during this “catch house” trial and predicts a “house” label, that would indicate that the brain currently encodes the house stimulus (stimulus N), purely as a result of an expectation of that stimulus, and that this representation persists even though the expectation has been violated (SI Appendix, Fig. S2C). If instead the stim-classifier predicts a face label, that would indicate a lingering prediction of the previous stimulus (N-1), for example because of left-over activity from the previous trial in the sequence. Similarly, a prediction of a phone label on this same “catch house” trial would indicate that the neural activity pattern was already representing the future stimulus N+1, even though the sequence is currently at stimulus N (SI Appendix, Fig. S2C). Note that these representations are not mutually exclusive, but could occur at the same time, as a sort of “superposition” state.

This cross-generalization approach was applied to the ISI periods and the catch trials. The example above describes the procedure for catch trials, but a similar reasoning can also be applied to the ISI periods. Note that in the case of ISI periods, a prediction of “current” refers to the next upcoming stimulus (stimulus N) after the present ISI period, and a prediction of “next” refers to the stimulus that is two stimuli away from the present ISI period (i.e., stimulus N+1). The cross-temporal generalization procedure was repeated separately for ISI and catch trials, and the proportion of predictions for the previous, current and next stimuli was counted. The null hypothesis in these analyses was that no relation existed between patterns of stimulus-evoked activity found in stim trials and patterns of activity recorded in the ISI periods and catch trials. Conversely, above-chance decoding of the “stim classifier” on the ISI and/or catch trials would mean that the ISI and/or catch trials carried sensory-like stimulus information even when no sensory input was available, purely as a result of top-down expectations.

Using this approach, for each sequence position (i.e. previous, current, next with respect to each trial onset) and period of interest (i.e. ISI period of all trials, and post-trial onset of catch trials), we tested whether there were significant above chance predictions of stimulus identity (Fig. 3). In the ISI period, the “current” upcoming event (stimulus N from the example above) was predicted significantly above chance (cluster test, P = 0.0001, Fig. 3B). This result indicates that stimulus-specific patterns of activity evoked by stimulus presentation on stim trials were already found in the ISI period, suggesting the emergence of a sensory template of the expected upcoming stimulus. No significant proportion of predictions for the previous sequence position was found (Fig. 3A), suggesting that stimulus-evoked patterns of neural activity of the preceding stimulus in the sequence were not present anymore in the ISI period. For the next sequence position, i.e. stimulus N+1 which was two stimuli away, one cluster was marginally significant toward the end of the pre-trial onset period (cluster test, p = 0.0534, Fig. 3C). To investigate the temporal dynamics of the reinstatement of stimulus-evoked neural patterns we computed the average classification accuracy across training time points (i.e, the Y-axis in A-C). The time course of the current sequence position (averaging across the Y-axis of Fig. 3B) exhibited a temporal cluster of significantly above chance predictions (P = 0.0001, Fig. 3G, blue curve). The time course of the next sequence position (averaging across the Y-axis of Fig. 3C) exhibited a smaller and later temporal cluster of significantly above chance predictions (P = 0.0128 Fig. 3G, red curve).

**Figure 3.**
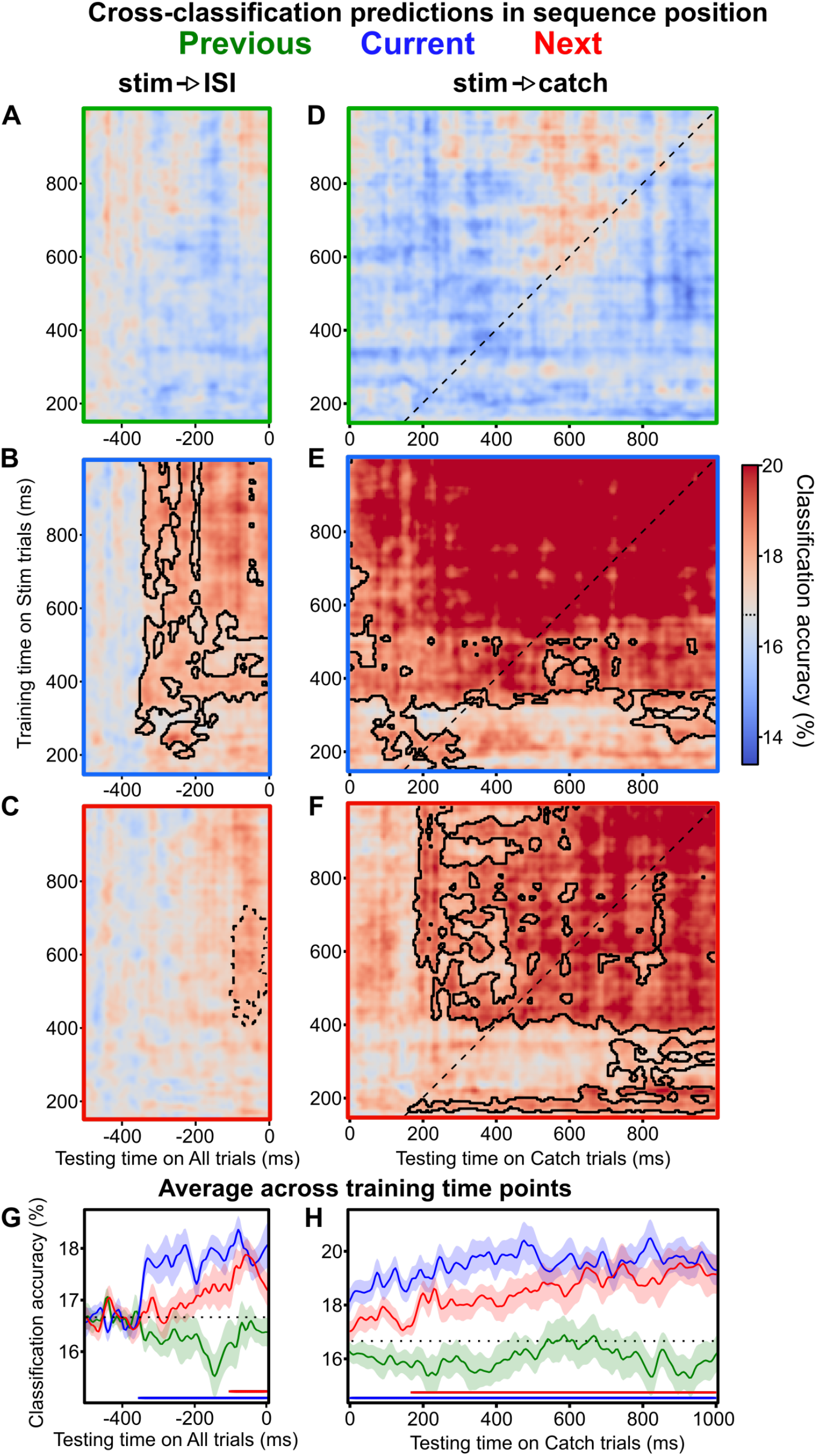
Decoding of ISI and catch trials using stimulus-evoked activity. A stimulus classifier was trained to discriminate between the six stimuli on stim trials. It was then used to predict stim identity on all ISI periods (A, B, C, G), and catch trials (D, E, F, H). We separately examined the time course of decoding performance for predictions of the preceding, current or next stimuli of the sequence (with respect to each trial onset; green curves represent the prediction of the previous stimulus in the sequence (N-1), blue curves represent the prediction of the current stimulus (N) and red curves represent the prediction of the next stimulus (N+1)). **A-C**. Cross-temporal prediction on all trials in the ISI period (−500 to 0ms prior to trial onset). A classifier was trained at time points from 150 to 1000ms of stim trials (vertical axis: training time on stim trials), and tested on all time points of the ISI period (horizontal axis: testing time in ISI period). Black contours indicate clusters of significant above-chance proportion of predictions (panel B: P < 0.0001). Black dashed contours indicate a marginally significant cluster (panel F: P = 0.0534). **D-F**. Cross-temporal prediction on catch trials. The same stim classifier was tested on catch trials (0 to 1000ms relative to catch trial onset). The black dashed line represents the diagonal of the temporal generalization matrix, i.e. classifiers trained and tested on the same time point in stim and catch trials. Black contours indicate clusters of significant above-chance proportion of predictions (cluster test, panel E: P = 0.0001, panel F: P < 0.001). **G-H.** Average of the prediction proportions across training time points (i.e., average along the Y-axis of the panels above). The black dotted line indicates chance-level (1/6). The horizontal lines below the curves indicate clusters of significant above-chance predictions. In panel G, the current (blue curve, P < 0.0001) and next (red curve, P = 0.0128) predictions were significantly above chance. In panel H, the current (blue curve, P < 0.0001) and next (blue curve, P < 0.002) predictions were significantly above chance.

Decoding performance on catch trials also indicated that sensory-like stimulus templates were established when the stimuli were expected, but not shown. On these catch trials, the current expected event (i.e., the omitted stimulus N from our example above) and the next event in the sequence (i.e., stimulus N+1), were predicted significantly above chance, on average across trials (cluster test, P = 0.0001, Fig. 3E and P < 0.001, Fig. 3F respectively). No significant proportion of predictions for the previous sequence position was found (Fig. 3D). To investigate the temporal dynamics of the reinstatement of stimulus-evoked neural patterns in catch trials we computed the average decoding performance across training time points (Fig. 3H). The time courses exhibited the same pattern of results: current sequence position (N) and next sequence position (N+1) were found to be significantly above chance (cluster test, P = 0.0001 and P < 0.002, respectively for current and next predictions). The time course of the N+1 stimulus prediction started later in the trial time course (approximately 200ms later) than the current stimulus N prediction (Fig. 3H). These results show that the brain maintains the identity of the current expected stimulus even when it is omitted, and after a short delay, starts encoding the next upcoming stimulus of the sequence.

Thus, during sequence learning, reinstatement of stimulus-evoked neural representations take place prior to expected stimulus onset (ISI periods) as well as when a stimulus is omitted in the sequence (catch trials). The brain starts representing future stimuli several hundred milliseconds before they actually appear, and critically, this stimulus representation is not erased when incoming sensory information indicates that the stimulus has not occurred. Instead, the brain actively represents information about the expected but omitted stimulus, and starts predicting the next stimulus to come, despite disruptions of the expected events.

### Spectral profile of predictive patterns of neural activity

Finally, to characterize the neural mechanisms supporting reinstatement of stimulus representations purely by top-down mechanisms, in the absence of bottom-up sensory input (i.e. catch trials), we tested whether specific oscillatory components exhibited selective patterns of activity. We performed a time-frequency decomposition of single catch trials in the post-trial onset period and used a classifier at each time and frequency point to decode the expected event. This analysis yielded a map of classification accuracy at each time point and frequency (Fig. 4A). No significant clusters were found in these maps using cluster-based permutations (P > 0.16), possibly due to the large number of multiple comparisons. To test whether certain frequencies carried information about the expected event, we averaged classification accuracies in these time-frequency maps across time and performed a cluster-based permutation analysis. We found a significant cluster of above-chance classification accuracies in the alpha-beta range (P < 0.05, Fig. 4B). This result suggests that neural activity selective to the expected stimulus was present in the alpha-beta frequency bands.

**Figure 4.**
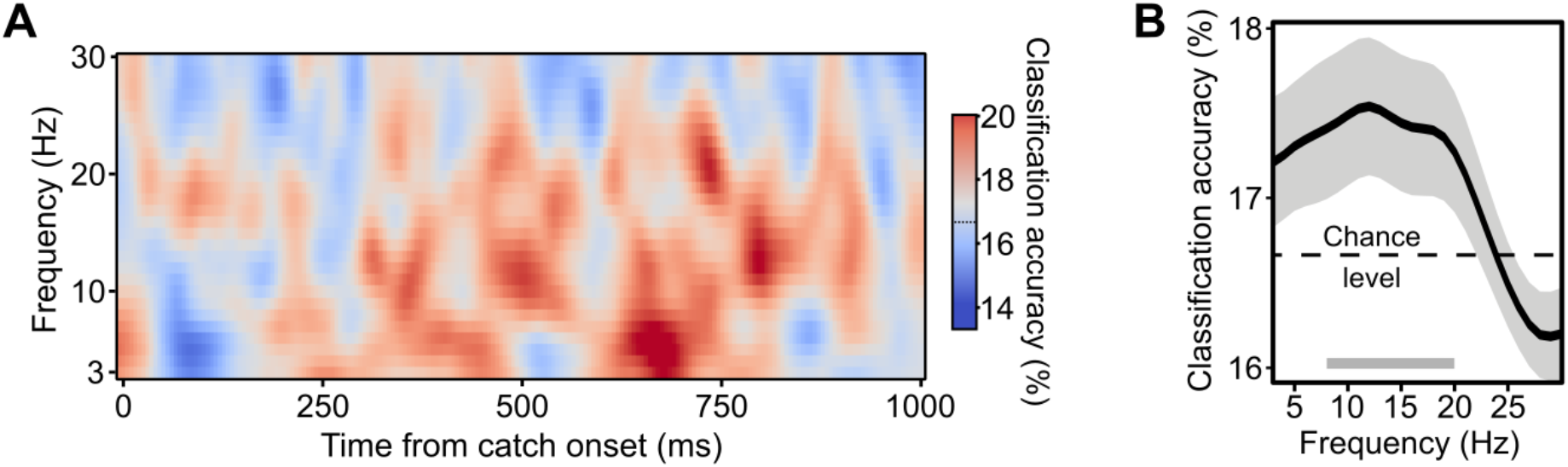
Spectral components of predictive neural patterns. **A.** Time-frequency map of classification accuracies. **B.** Frequency profile of classification accuracy (average accuracy over time). Gray bar at the bottom represents the significant cluster of classifier accuracy going from 9Hz to 20Hz.

## Discussion

In this study we investigated whether visual sequence learning induced anticipatory reinstatement of expected stimulus neural representations. We found that (1) both prior to expected stimulus onset and when the expected stimulus was omitted, EEG activity exhibited neural patterns selective to the expected stimulus. (2) The selective patterns of neural activity present prior to the onset of the expected stimulus generalized to periods in which the expected stimulus was omitted, and (3) following this reinstatement of the representation of the omitted stimulus, patterns selective to the next stimulus in the sequence emerged, suggesting a proactive mechanism forming predictions about future events. (4) Finally, these selective patterns were mostly evoked in the topography of alpha-beta bands oscillations, confirming the role of these oscillatory bands in carrying feedback signals across cortical areas (Spitzer and Haegens, 2017; Alamia and VanRullen, 2019). These results suggest that the brain actively reinstates representations for upcoming stimuli in a predictable sequence, and that these patterns of activity persist even when the expected event eventually fails to occur. This mechanism might play an important role in maintaining sequence order, in the face of eventual disruptions.

An important result of our study is the finding that during catch trials, neural activity patterns simultaneously represented the current expected stimulus, as well as the next stimulus in the sequence (Fig. 3). This finding suggests an active mechanism for predicting future events. This decoding of stimulus-selective neural patterns during the catch trials relied on using the “stim” trials as the template for sensory representations. However, since the stim trials themselves were not free of expectations (the subjects were performing the task and anticipating the upcoming stimuli), we cannot exclude the fact that the representations reinstated during the catch trials did not also partly include a reinstatement of the expectation signal. It would be interesting in future research to test whether purely perceptual representations, allows for stimulus-selective decoding during catch trials.

Our study builds on and extends previous work on how expectations shape neural representations in anticipation of future events. In this study we focused on the anticipatory reinstatement of stimulus representations during the maintenance of an ongoing sequence of events. Previous studies have shown that expectations can bias perception (Chalk et al., 2010; Sotiropoulos et al., 2011; Pajani et al., 2015), improve perceptual performance (Kok et al., 2012, 2014; Rohenkohl et al., 2012; Wyart et al., 2012) and action preparation and execution (Nobre et al., 2007). Numerous studies have explored the neural bases of expectation signals (Kok et al., 2017; Blom et al., 2020; Boettcher et al., 2020). For instance, it has been shown that expectations sharpen and bias neural representations of expected features in early visual cortex (Kok et al., 2013, 2014). A recent magnetoencephalography (MEG) study revealed that expectations create sensory templates before the expected stimulus presentation and that this template influenced post-stimulus neural representations (Kok et al., 2017). Another recent study, similarly showed that a visual object’s spatial movement is anticipated in a similar fashion, by reactivating neural representations of this expected stimulus position (Blom et al., 2020). Our results complement these studies by demonstrating that reinstatement induced by expectations of future events takes place in more complex settings (i.e. learning a sequence of stimuli instead of single stimulus association pairs), and that multiple future events can be simultaneously represented in EEG activity patterns.

Our findings resemble forward prediction found in navigation, and more generally sequence-learning, found in medial temporal lobe structures (Johnson and Redish, 2007; Davachi and DuBrow, 2015; Reddy et al., 2015, 2019). Indeed, learning sequential event order allows one to form predictions and adequately prepare future actions. This ability has been shown to critically depend on intact hippocampal circuits and that sequence learning strengthens relational representations in this region (Schapiro et al., 2012; Reddy et al., 2015). To date it has been unclear how sequence learning affects distinct neural representations of expected stimuli, prior to their expected apparition or in their absence. In this study, we show that neural representation of expected stimuli embedded in a sequence are evoked prior to expected onset, but also that a proactive reinstatement of future stimuli takes place. Furthermore, we found selective activity patterns evoked in the alpha-beta band which is in line with recent studies showing increased alpha-beta synchrony between hippocampal and prefrontal cortices (Brincat and Miller, 2015), and increases in parieto-occipital alpha-beta amplitude related to future events in learned sequences (Crivelli-Decker et al., 2018).

We found evidence that alpha-beta band amplitude topography was selective to the expected stimulus. This is in line with accumulating evidence from experimental and modeling work pointing towards a critical role of alpha band oscillations in instantiating perceptual priors (Sherman et al., 2016; Chang et al., 2017) and reflecting predictive coding computations (Han and VanRullen, 2017; Alamia and VanRullen, 2019). Furthermore, it has recently been shown that shared neural representations of visual objects in perception and imagery are expressed in the alpha band (Xie et al., 2020), which suggests a functional overlap between the reinstatement of expected stimulus representations observed in our study, and mental imagery.

Beta band oscillations, on the other hand, have been shown to provide feedback signals across cortical areas (Bastos et al., 2012; Michalareas et al., 2016; Richter et al., 2017), and to support top-down processes such as attentional preparation and reinstatement of task-relevant content (Engel and Fries, 2010; Spitzer and Haegens, 2017). The involvement of beta oscillations in expectation processes is not restricted to the visual domain, e.g. in the auditory domain (Todorovic et al., 2015), which suggest a domain general mechanism carrying expectation signals in the beta band.

Taken together these results suggest an important role of alpha and beta oscillations in top-down processes, in allowing synchronization between sensory expectations and an external stimulation rhythm (Arnal et al., 2015; Todorovic et al., 2015; Chang et al., 2017) and thereby, carrying information about the nature of expected stimulus (Bastos et al., 2012; Michalareas et al., 2016; Richter et al., 2017; Spitzer and Haegens, 2017). Our results are thus congruent with this proposed role of alpha-beta oscillations, i.e. in our study representing expectations of the next stimulus to come in the sequence. The range of frequencies supporting these expectation signals seem to vary across the alpha and beta bands. Recent studies have shown that task demands affect the specific frequency of neural oscillations (Wutz et al., 2018; Senoussi et al., 2020). It would therefore be of interest in future research to investigate whether task properties, such as sensory modality, task difficulty, or the nature of the task-relevant expectations (e.g. stimulus identity, timing) differentially recruit specific frequency bands.

Our results also link the fields of sensory expectations, predictive coding and sequence learning (Davachi and DuBrow, 2015; de Lange et al., 2018) by providing specific mechanisms supporting the reinstatement by the MTL of expected events. A large body of literature has shown that theta oscillations are a core mechanism supporting short-term memory and more specifically the ordinal organization of memoranda (Fuentemilla et al., 2010; Davachi and DuBrow, 2015; Crivelli-Decker et al., 2018; Peters et al., 2018; Reddy et al., 2019). It will therefore be an important avenue for future research to investigate how these alpha-beta oscillations are generated, how they carry selective information about expected events, and the interaction with MTL theta oscillations (e.g. through cross-frequency coupling).

To conclude, using MVPA and time-frequency decomposition we found that visual sequence learning elicits selective patterns of EEG activity prior to expected event onset and in the absence of the expected stimulus. These patterns were similar to stimulus evoked activity, and patterns selective to the next-to-come event emerged towards the end of an event period, suggesting a constant prediction of future sensory events. Finally, selective neural activity was found to be expressed in the alpha-beta band, which has been associated with feedback signals and sensory expectation, confirming its essential role in providing task-relevant contextual information for efficient processing of future events. These results provide important insights into how the brain implements predictive mechanisms for future when expectations are violated, based on regularities learned in the past.

## Materials and Methods

### Participants and stimuli

Sixteen participants (age 21-45 years, five females) were recruited in this experiment. All participants had normal or corrected-to-normal vision and no history of neurological problems. One participant was excluded due to chance level behavioral performance during test trials in half of the blocks. All participants provided written informed consent and received monetary compensation for their participation. The study was approved by the local ethics committee “Sud-Ouest et Outre-Mer I” and followed the Code of Ethics of the World Medical Association (Declaration of Helsinki).

Six stimuli from six different object categories, i.e. face, camel, car, house, pine cone and phone, were gathered from internet. The selection of visual object categories was based on previous studies which computed the distance in neural representations between exemplars from different visual object categories (Kriegeskorte et al., 2008; Carlson et al., 2013). The rationale was to maximize the distance in the multivariate EEG response between stimuli to maximize their discriminability. All stimuli were equalized in the 2D Fourier power spectrum in order to diminish low-level confounds in the stimuli set.

### Experimental procedure

Participants were instructed to learn a stimulus sequence of six images (see Fig. 1A), e.g. pine cone -> camel -> face -> house -> phone -> car. The images in the sequence were presented in a fixed, pre-determined order, and the sequence was repeated 480 times in total across the whole experiment. The stimulus order was randomized for each participant. Participants were seated 60cm from the screen. Each image was presented for 1s preceded and followed by an inter-stimulus interval (ISI) of 0.5s.

To further the impression of a sequence of images we used the following display arrangement (SI Appendix, Fig. S1A): Each image was presented at the center of the screen while placeholders (empty black squares) were presented to the left and right of the central image. At the end of the 1s presentation period, the central image was replaced by a black placeholder and all the placeholders moved one “step” forward in a clockwise direction for the duration of the ISI, such that each placeholder eventually occupied the next placeholder position. At the end of the ISI, the placeholder that now occupied the central position was replaced by an image. The viewer’s subjective impression at the end of the ISI interval was that the central image had been hidden, and then moved clockwise, while the central position was replaced by the next image. The central placeholder was at the center of the screen, and covered 6° of visual angle.

On each trial of the sequence, one of three possible types of trial types could appear (Fig. 1B): 1) the expected stimulus from the sequence was displayed for 1s (stim trials, 45% of all trials), 2) the expected stimulus was not shown, but was replaced by a gray square for 1s (catch trials, 45% of all trials), or 3) test trials (10% of trials), in which, instead of showing the expected stimulus in the sequence, two choice stimuli were displayed and participants had to report which of the two was the expected image. The test trials were self-paced. Participants were told that they could rest their eyes during test trials, and respond to the test trial when they were ready to resume the sequence. No feedback was provided on test trials.

The experiment consisted of eight blocks, each lasting approximately 10 minutes. Stim, catch and test trials were randomly interleaved. Each block contained 60 loops of the six-image sequence, resulting in 360 trials per block and 2880 trials in total for each participant. This design resulted in 288 test trials (10% of the total), and 1296 stim and 1296 catch trials (45% each of the total).

A green fixation point was present at the center of the screen at all times and participants were instructed to fixate this dot during the whole experiment except for test trials during which they could move their eyes freely. In order to familiarize participants with the task and experiment layout, they were shown 10 trials (stim and catch trials), of another sequence than the one they had to learn, prior to the start of the experiment.

### EEG acquisition and preprocessing

64-channel EEG was recorded using a BioSemi Active-Two system at a sampling rate of 1024 Hz. A three-channel EOG was also recorded to monitor horizontal and vertical eye movements and blinks. Data were downsampled offline to 200Hz and notch-filtered at 50Hz to remove electrical line noise. We defined trial start as the onset of the presentation of a stimulus (on stim trials), the gray square (on catch trials), or the two choice stimuli (test trials). We extracted epochs from 500ms before trial onset to 1000ms post trial onset, yielding epochs of 1500ms, in which 0ms represents the trial onset. Raw EEG time-courses were screened manually on a trial-by-trial basis to reject visible artifacts, eye movements or blinks. Baseline correction was applied by subtracting the average activity from −500 to −400ms relative to stimulus onset, for each electrode and trial independently. The median number of rejected trials (for non-test trials) across participants was 78/2592.

We use the term trial type to refer to whether a trial was a stim, catch or test trial, and the term events for the 6 possible events for each trial type, i.e. the images in the sequence. For stim trials there was therefore 6 stim events: stim-face, stim-car, stim-house, stim-camel, stim-pine cone and stim-phone. For catch trials there were 6 catch events depending on which image should have appeared but was replaced by a gray square: catch-face, catch-car, catch-house, catch-camel, catch-pine cone and catch-phone. And for ISI periods there 6 ISI events depending on which image was about to appear at the end of the ISI period: ISI-face, ISI-car, ISI-house, ISI-camel, ISI-pine cone and ISI-phone.

There was, on average across participants, 201 trials per for each event of stim and catch trials (i.e. 6 stim events, 201 trials each; 6 catch events, 201 trials each). ISI periods preceding stim, catch, and test trials were pooled because the ISI periods were not predictive of the subsequent trial type. Therefore, on average, for each event, an additional 42 ISI periods preceding test trials were added to the ISI periods preceding stim and catch trials. There was thus, on average across participants, 243 trials per for each ISI event.

### Time-frequency decomposition

We computed time-frequency amplitude through a complex Morlet wavelet decomposition on single trials implemented in the MNE-Python suite (Gramfort et al., 2013, 2014). The frequencies ranged from 3 to 30hz (28 linearly spaced steps) with the number of cycles linearly increasing from 1.5 to 5 cycles.

### EEG pattern classification

Our goal was to investigate whether selective patterns of neural activity to the expected stimulus arise 1) prior to trial onset (i.e. during the ISI period) and 2) in the absence of the expected stimulus (i.e. catch trials). To test whether selective activity patterns were elicited, we used multivariate pattern analysis (MVPA) on scalp topographies at each time point to classify sequence information in the ISI periods, the stim and the catch trials.

### Classification analysis settings

In all analyses, we performed time-resolved 6-way (e.g. discriminate the 6 events from each other) classification analyses on scalp topographies including all electrodes. These analyses were carried out on raw electrical potentials (i.e. broadband EEG signal after preprocessing) and time-frequency amplitudes. For all classification analyses we trained and tested a linear discriminant analysis (LDA) classifier using a least-square solver combined with automatic shrinkage using the Ledoit-Wolf lemma. The 6-way classification was performed using a one-versus-all multi-class procedure. We used a Stratified 5-fold cross-validation procedure. Both classifier and cross-validation were implemented using the Scikit-learn Python toolbox version 0.21.2 (Pedregosa et al., 2011).

For all classification results we convolved the classification accuracy time-courses with a gaussian kernel, with parameters μ=0 and □=1, to lessen classification accuracies noise. The gaussian kernel was trimmed at 4*□, meaning that for each time point, three time points preceding, and following it were affected by the convolution, i.e. a 35ms temporal window.

### Pseudo-trials for classification

To increase signal-to-noise ratio, in all analyses we averaged single trials in order to make 5 pseudo-trials for each of the six stim, catch and ISI events (Isik et al., 2014; Grootswagers et al., 2016). The classification analysis was performed on the pseudo-trials. The pseudo-trials were made separately for each of the 6 events of ISI periods, stim and catch trials. Each classification section below describes the number of trials that were used to make the pseudotrials. Because each pseudo-trial was constructed from an arbitrary selection of single trials, we repeated the pseudo-trials creation 50 times (each with a different and random selection of single trials) to minimize sampling bias, and re-ran the classification analysis. We averaged decoding results across re-samples in all analyses.

### Preceding trial type control

Because of the temporal proximity between stimulus and catch trials, i.e. ISI of 500ms, we carried out a control analysis to investigate whether the preceding trial type affected classification accuracies. That is, is decoding performance on a particular trial N biased by a leakage of stimulus information from the preceding trial (trial N-1)?

We trained a classifier on training data at time point *t* and tested it on test data at time point *t*. This procedure was repeated for each time point, yielding a 1-dimensional array of classification accuracies. This analysis allows to estimate whether there is a difference in topography patterns between classes, as a function of time. For each trial N, we performed temporal classification at each time point. This was done separately for the stim and catch trials. Because, we sought to estimate the time at which the preceding trial-type (trial N-1) did not significantly affect the classification accuracy of the current trial (trial N) we used a Threshold-Free Cluster Enhancement (TFCE) procedure (start = 0, step size = 0.1, 10,000 permutations). TFCE creates a threshold for each data point under the null hypothesis, hence making inference at the data point level, and not at the cluster level as in fixed-threshold cluster-based permutation tests (Maris and Oostenveld, 2007; Smith and Nichols, 2009; Sassenhagen and Draschkow, 2019), which allowed us to estimate the latest time point at which the preceding trial type (N-1) affected classification accuracy of the current trial (N).

Using the TFCE procedure we tested whether there was a difference in decoding performance depending on whether trial N was preceded by a stim or catch trial (i.e., we test the influence of trial N-1 on trial N). We found significantly higher classification accuracies on trial N when it was preceded by a stim trial, compared to when it was preceded by a catch trial, indicating residual stimulus activity that persisted through the ISI period between trials (P < 0.05 corrected for multiple comparisons, see SI Appendix, Fig. S3A). On the Nth stim trial, the influence of the preceding (N-1) stim trial was observed as early as −385ms and until 85ms after trial onset. For the Nth catch trial, the effect of the preceding (N-1) stim trial was observed as early as −265ms and until 230ms after trial onset.

To ensure that leakage of information from previous trials did not persist even longer in time, we performed another control analysis in which we accounted for the influence of trial N-2 on trial N. In this analysis, we first selected the subset of trials that were preceded by a catch trial (i.e., trial N-1 is a catch trial). This subset of trials is thus not biased by stimulus information from the preceding trial. We then tested the influence of trial N-2 on the current trial N. Is there a difference in decoding performance on these trials N depending on whether trial N-2 is a stim or catch trial? No difference was found in decoding performance on trial N when N-2 was a stim trial vs. a catch trial. This was true both when N itself was a stim trial or a catch trial. These results thus indicate that trial N-2 did not influence classification accuracies on trial N.

Thus, for temporal generalization decoding analyses (see below) we only included trials (N) that were preceded by a catch trial (i.e., N-1 was a catch trial). For classification on stim and catch trials, this selection reduced the average number of trials to 112, on average, for each of the six stim event and each of the six catch events. For ISI periods, on average an additional 23 ISI periods preceding test trials were added to the ISI periods preceding stim and catch trials. There were thus 247 trials on average for each of the six ISI events, i.e. 112 ISI periods preceding stim trials, 112 ISI periods preceding catch trials and 23 ISI periods preceding test trials.

In the cross-temporal generalization analysis (see below), a classifier was trained only on stim trials (stim-classifier) to discriminate between the six stimulus events. The time window for training the classifier (150-1000ms post trial onset) was chosen to only include time points that did not show an influence of the preceding N-1 trial (the last time point that showed a difference in decoding performance on stim trial N depending on whether N-1 was a stim or catch trial was 85ms post trial onset). Thus, this classifier learns to discriminate between the six visual stimuli, and was tested on the ISI periods and the catch trials to determine whether sensory stimulus activation patterns were found even in these time periods when no stimulus was present.

### Classification analysis: temporal generalization (Fig. 2A-B, and SI Appendix, Fig. S1B)

In the temporal generalization analysis, we trained a classifier on the training data at time point *t*, and tested it on test data at every time point *t*’, including the time point *t*’=*t*. This procedure was repeated for each training time point *t*, yielding a 2-dimensional map of classification accuracies of training time vs. testing time (Fig. 2). This analysis allowed us to estimate whether a discriminative pattern appearing at a certain time point was informative for classification at other time points in the trial, e.g. because this neural activity pattern was sustained or re-appeared at different moments of the trial (King and Dehaene, 2014).

As explained in the section “Preceding trial type control” above, only trials preceded by a catch trial (N-1 catch) were used in temporal generalization analyses. Thus, for temporal generalization analyses on stim and catch trials, the average number of trials per event and per trial type (stim or catch) was 112. Therefore, for each event (e.g. catch-face), we averaged 22 single trials (on average across participants) to make 5 pseudo-trials per event.

For temporal generalization analyses on ISI periods the average number of trials per event was 135. Therefore, for each event (e.g. ISI-face), we averaged 49 single trials (on average across participants) to make 5 pseudo-trials per event.

We then used a Stratified 5-fold cross-validation procedure, in which a classifier was trained on 4 pseudo-trials per event x 6 events = 24 pseudo-trials for training, and tested on the remaining 1 pseudo-trial per event x 6 events = 6 pseudo-trials for testing.

### Classification analysis: cross-temporal generalization from ISI periods to catch trials (Fig. 2C)

In the cross-temporal generalization we performed the same procedure as in temporal generalization but the classifier was trained and tested on independent sets of data. Two cross-temporal generalization analyses were performed. In the first analysis (Figure 2C), a 6-way classifier was trained to discriminate between the ISI periods, and tested on catch trials. The ISI intervals were split into 6 groups, depending on which event they preceded. More precisely, the classifier was trained on ISI data at time point *t* and tested at time points *t*’ of the catch trials. This cross-decoding method allowed us to test whether selective patterns of neural activity generalize across conditions (Reddy et al., 2010; Kaplan et al., 2015; Senoussi et al., 2016): do brain patterns elicited in expectation of a given stimulus (in the ISI period) correspond to brain patterns that are recorded when the expected stimulus is not shown (on catch trials)?

As explained in the section “Preceding trial type control” above, only trials preceded by a catch trial (N-1 catch) were used in this analysis. We used Stratified 5-fold cross-validation on ISI periods preceding catch trials in the training set and catch trials in the testing set. This was done to prevent having the ISI period preceding a specific catch trial and the post-trial onset period of this trial (i.e. the catch trial) being used respectively as training and testing data which would result in spuriously high decoding accuracy, i.e. double-dipping (Kriegeskorte et al., 2009).

Therefore, for one cross-validation fold we used all ISI periods preceding stim trials, all ISI periods preceding test trials, but only 4/5^th^ of the ISI periods preceding catch trials. The remaining 1/5^th^ of ISI periods preceding catch trials was left-out in this cross-validation fold. The catch trials, i.e. the post-trial onset period following the left-out ISI periods) were used as the test dataset in this cross-validation fold. This selection procedure was repeated five times in order to use all ISI periods preceding catch trials in the training dataset and all catch trials in the testing dataset.

Therefore, for each event (e.g. ISI-face) and each cross-validation fold the training dataset was composed of 112 (all ISI preceding stim trials) + 90 (ISI preceding catch trials, i.e. 4/5^th^ of the total 112 ISI periods preceding catch trials) + 23 (all ISI preceding test trials) = 225 single trials (on average across participants). For the testing dataset, for each event (e.g. catch-face) and each cross-validation fold, 1/5^th^ of all catch trials were used, which amounts to 22 trials.

As for other classification analyses, we created 5 pseudo-trials per event, separately for the training and testing datasets. For the training dataset we therefore averaged 45 trials per ISI event to create one pseudo-trial. For the testing dataset we therefore averaged 4 trials per catch event to create one pseudo-trial.

The classifier was then trained on all 5 pseudo-trials per ISI event x 6 ISI events = 30 pseudo-trials for training, and tested on the 5 pseudo-trial per catch event x 6 catch events = 30 pseudo-trials for testing.

### Classification analysis: cross-temporal generalization from stim to ISI and catch trials (Fig. 3)

In the second cross-temporal generalization analysis (Fig. 3), a 6-way classifier was trained on only the stim trials (stim-classifier), to discriminate between the six images, at each time point *t*. It was then tested in two separate analyses on 1) time points *t*’ of the ISI periods and 2) *t*” of the catch trials. This analysis allowed us to determine whether sensory stimulus information recorded when participants were viewing the stimuli corresponded to brain patterns induced by purely top-down expectations of these stimuli (in the absence of sensory input). The section “Preceding trial type control” above explains which time points of stim trials were used for training the stim classifier.

This procedure was repeated for each training time point of the training dataset and it yielded six 2-dimensional maps of classifier predictions (training vs. testing time) for each analysis (i.e. on ISI periods and on catch trials). For instance, for the analysis in which we tested the stim-classifier on ISI periods (Fig. 3A-C, G), for one training time point *t* and one testing time point *t*’, we obtained six values representing the proportion of the six stim-events that were predicted (e.g. 14% of stim-car event, 18% of stim-house event, 29% of stim-face event, etc.). Across all training and testing time points these predictions thus formed six maps for the analysis in which we tested the stim-classifier on ISI periods.

The stim-classifier made a prediction about stimulus identity on the testing data (SI Appendix, Fig. S2), and on each test trial it could make a prediction of any of the six stimulus events. On each trial of the testing data, we counted the proportion of times the stim-classifier predicted the current stimulus, and the stimuli preceding and following it in the sequence. For example, in a sequence segment consisting of three stimuli: face-house-phone (SI Appendix, Fig. S2B), when testing the stim classifier on a house catch trial (or an ISI preceding a house trial), a prediction of “house” by the stim classifier would be counted as a prediction of the current stimulus. A prediction of “face” or “phone” would be counted as predictions of the preceding and following stimuli, respectively. This measure allowed us to determine whether brain activity patterns during the ISI and catch periods encoded stimulus identity, and if so, which stimuli in the sequence were being encoded. The null hypothesis was that no sensory-like stimulus information was present in EEG topographies of ISI periods and catch trials, hence each stimulus identity would be predicted at chance level (1/6 = 16.66%).

### Statistical analyses

Inferential claims about all classification analyses were based on cluster-based permutation test (Maris and Oostenveld, 2007) and reported following the recommendation of Sassenhagen & Draschkow (2019). Cluster-based permutation analyses assess the probability of observing a certain classification accuracy cluster size using a null distribution of cluster sizes generated through permutations. The clusters are made by temporally or spectrally adjacent samples exceeding a threshold defined using the Student T-value distribution by comparing classification accuracies to chance level (1/6 = 16%). This procedure yields a P-value corresponding to the probability of observing a cluster of classification accuracies exceeding the threshold based on a null distribution of cluster sizes while correcting for multiple comparisons.

For cluster-based permutation tests we used the *permutation_cluster_1samp_test* of the MNE toolbox v.0.18 (Gramfort et al., 2014), with a Hat variance adjustment, with parameter □=10^-3^, to correct for small pixel variances (Ridgway et al., 2012). Specifically, the procedure randomly permuted the sign of centered classification accuracy values, i.e. classification accuracies minus chance level (1/6), in a random subset of participants, computed the size of clusters exceeding the threshold in these surrogate data, and repeated that procedure to obtain a null distribution of cluster sizes. This procedure was repeated 10,000 times for each analysis and yielded cluster P-values (the clusters being 1-or 2-dimensional, depending on the analysis). From these analyses, P-values of <0.05 are depicted as straight lines below classification accuracy curves (Fig. 3G-H and SI Appendix Fig. S3), or black contour lines in 2D maps of classification accuracies (Fig. 2A-C, 3A-F).

### Activation patterns

Because the interpretation of multivariate classifiers’ weights (i.e., the class-selective topographies) represents a mixture of signal of interest and noise, we computed activation patterns using the method developed in Haufe et al. (2014). This method allows to extract the discriminative topographies used by the classifier to distinguish the different classes. Because all our analyses consisted of a 6-way classification, six activation patterns were produced at each time step. We computed the Student T-value for each event (e.g. catch-face) across all participants, rectified these T-value topographies by computing their absolute value, and averaged the six topographies. This produced one topography for each training data set (i.e. stim trials, catch trials, and ISI periods), for each time point.

In order to compute a measure of the informativeness of these activation patterns, we averaged these topographies in consecutive 100ms temporal window and weighted these values by the decoding performance. Specifically, topographies obtained after averaging across the six events were averaged in windows of 100ms, scaled to a standard deviation of 1 across electrodes, and multiplied by the average T-value of decoding performance across participants for that same time window.

## Supplementary Information Appendix

**Figure S1.**
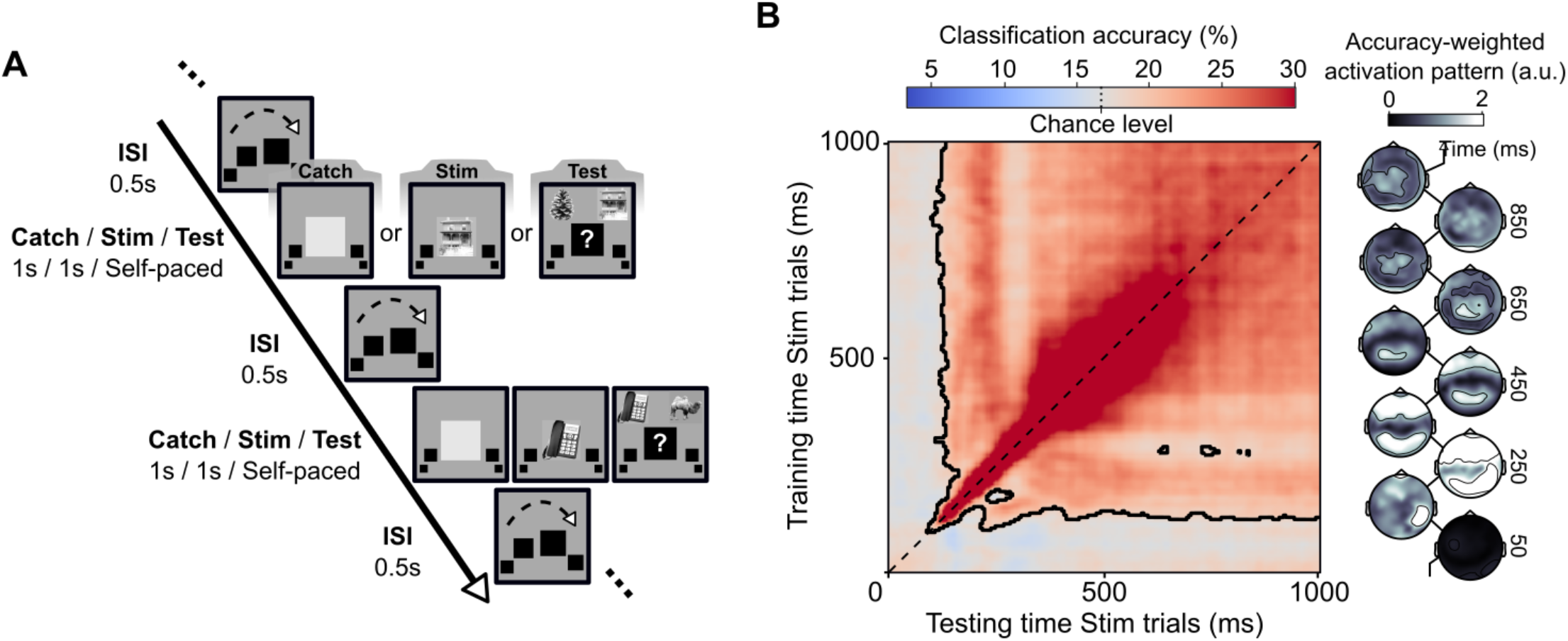
Trial structure and temporal generalization analysis on stim trials. **A.** Participants were asked to learn the order of a sequence of six stimuli. On each trial either one a catch, stim or test trials appeared. Each trial started with an inter-stimulus interval (ISI) of 0.5s followed by the presentation of an image (stimulus or gray square) for 1s, or a test trial during which participants had to report which of the two presented stimuli should have appeared in the sequence. During the image presentation period the image was presented at the center of the screen, flanked by black placeholders. During the ISI period, the placeholder turned black, replacing the central image, and all the placeholders moved in the clockwise direction (the dashed arrow in ISI screens is only to illustrate the movement and was not shown during the experiment). At the end of the ISI period, the central placeholder was replaced by the next image in the sequence. The trial type (stim, catch or test) order was randomized. **B.** A 6-way classifier was trained at each time point (vertical axis: training time) and tested on all time points (horizontal axis: testing time) to distinguish each stimulus event. The black dashed line represents the diagonal of the temporal generalization matrix, i.e. classifiers trained and tested on the same time point. Black contours indicate a cluster of significant above-chance classification accuracy (P = 0.0001). Topographies on the right side of the map represent activation patterns weighted by decoding performance (see Methods) for classifiers trained on stim trials.

**Figure S2.**
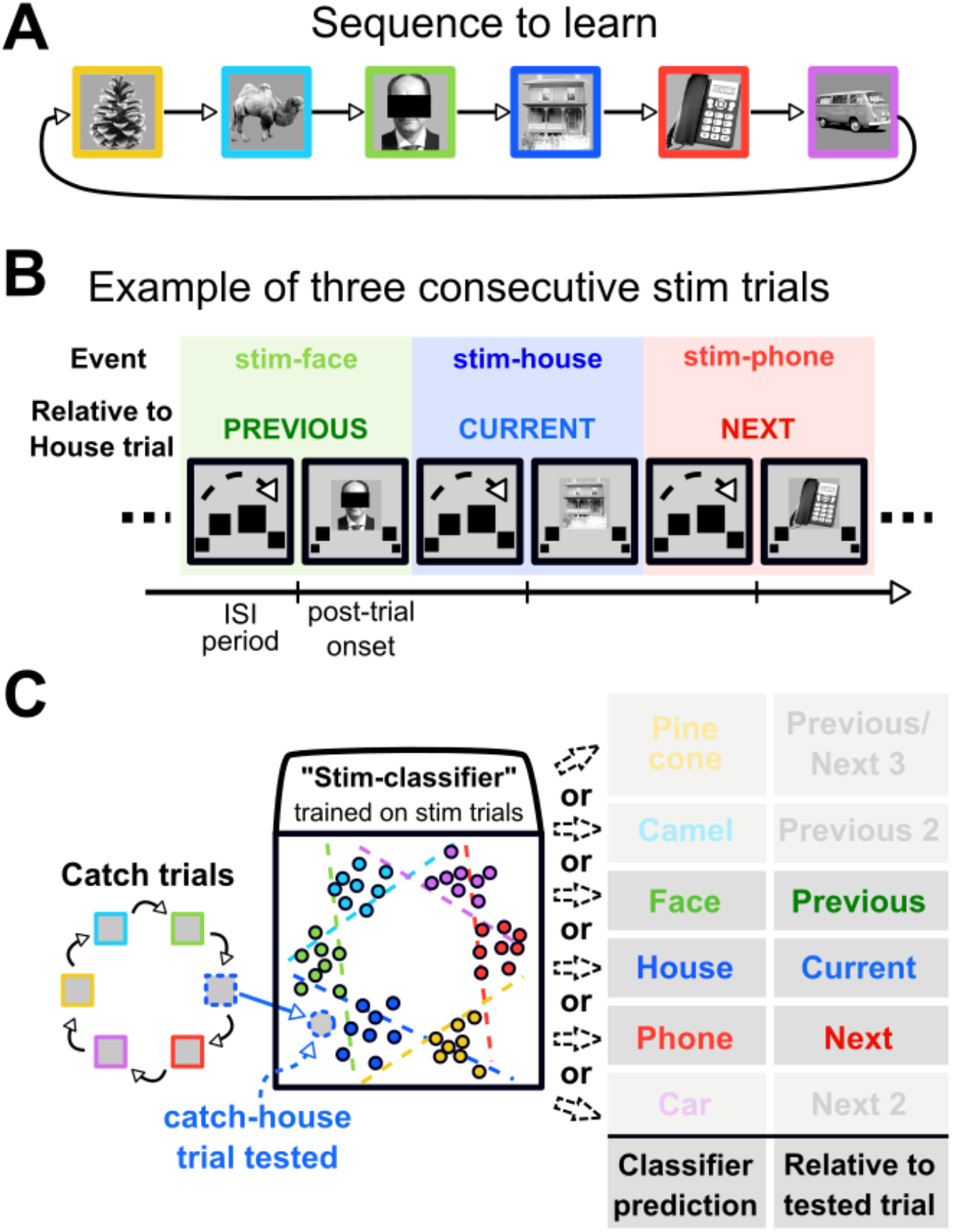
Cross-temporal generalization analysis. **A.** An example stimulus sequence to learn. **B.** A sequence segment showing three consecutive (stim) trials. Relative to the trial at the “House” position in the sequence, the “Face” trial is the previous one, and the “Phone” trial is the next one. **C.** A stim-classifier is tested on catch-house event data. The stim-classifier can predict any of the six stim events. If the classifier predicts a stim-house event, we considered it as a prediction of the “Current” position in the sequence. Alternatively, if the stim-classifier predicted a stim-face or a stim-phone, these are counted as predictions of the previous and next events in the sequence (and could arise for example by left over activity of the preceding stimulus, or anticipation of the next stimulus respectively).

**Figure S3.**
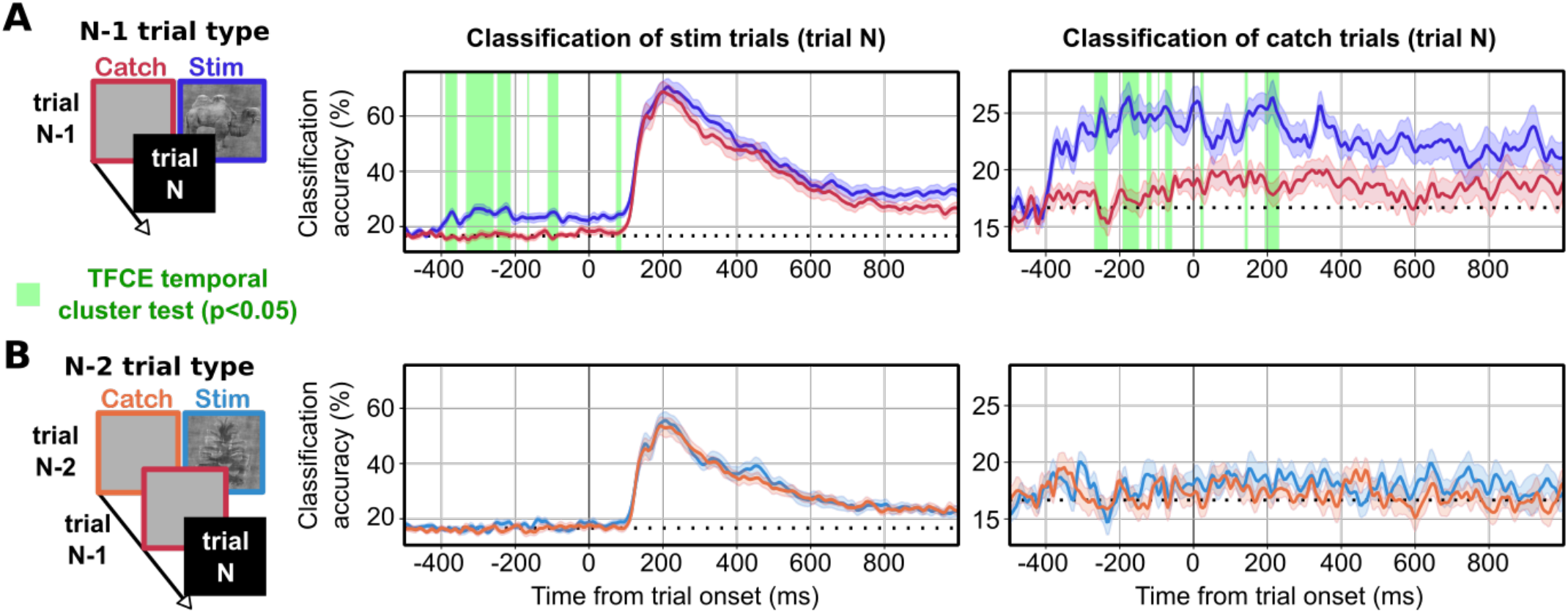
Preceding trial type analysis. **A.** Classification accuracies for classified trials (trial N) that were preceded by a stim or catch trial (N-1). The diagram on the left represents the color code: dark red when trial N-1 was a catch trial, dark blue when trial N-1 was a stim trial. Middle and right plots depict average temporal classification on trial N when trial N was a stim trial and a catch trial respectively. Classification accuracy was significantly higher in both the stim and catch trials (trial N) when the preceding trial (trial N-1) was a stim trial (blue curves) compared to when the preceding trial (trial N-1) was a catch (dark red curves), indicating residual stimulus activity from the preceding stim trial. **B.** In this analysis we selected the subset of trials (N) that were preceded by a catch trial. We then examined how decoding performance on trial N was influenced by whether trial N-2 was a stim or catch trial. The diagram on the left represents the color code: orange when trial N-2 was a catch trial, light blue for when trial N-2 was a stim trial. Middle and right plots depict temporal classification on trial N, when it was a stim and catch trial respectively. Note that there is no significant difference between trials in which N-2 was a stim or a catch trial for both trial N types (stim or catch), indicating that residual activity of the N-2 stimulus is not observed in trial N. In both plots, green areas represent a significant difference (p<0.05) computed between the two curves in each plot using a cluster-based permutation (TFCE procedure, see Methods). Colored shading around the curves indicates standard error of the mean across participants.

## Conflict of Interest

The authors declare no competing financial interests.

## Acknowledgments

This work was supported by the French Research Agency (Vis-Ex ANR-12-JSH2-0004) to L.R. and the European Research Council Grant (P-CYCLES 614244) to R.V.

## Code and data availability

Raw EEG data can be found on this OSF repository: https://osf.io/5ycfu/?view_only=d790a6a279e74e81868801b22c57fc75.

Behavioral data and scripts to reproduce all analyses and figures can be found on this Github repository: https://github.com/mehdisenoussi/assolearneeg

## Notes

https://github.com/mehdisenoussi/assolearneeg

### Competing Interest Statement

The authors have declared no competing interest.

